# Active inference through whiskers

**DOI:** 10.1101/2021.07.16.452665

**Authors:** Francesco Mannella, Federico Maggiore, Manuel Baltieri, Giovanni Pezzulo

## Abstract

Rodents use whisking to probe actively their environment and to locate objects in space, hence providing a paradigmatic biological example of active sensing. Numerous studies show that the control of whisking has anticipatory aspects. For example, rodents target their whisker protraction to the distance at which they expect objects, rather than just reacting fast to contacts with unexpected objects. Here we characterize the anticipatory control of whisking in rodents as an active inference process. In this perspective, the rodent is endowed with a prior belief that it will touch something at the end of the whisker protraction, and it continuously modulates its whisking amplitude to minimize (proprioceptive and somatosensory) prediction errors arising from an unexpected whisker-object contact, or from a lack of an expected contact. We will use the model to qualitatively reproduce key empirical findings about the ways rodents modulate their whisker amplitude during exploration and the scanning of (expected or unexpected) objects. Furthermore, we will discuss how the components of active inference model can in principle map to the neurobiological circuits of rodent whisking.

## 1. Introduction

Rodents use whisking to probe actively their environment and to locate objects in space - hence providing a paradigmatic biological example of *active sensing* [1, 2, 3]. Active whisking has been studied in many contexts, such as during the perception of object location [4, 5, 6, 7] and the discrimination of shapes [8].

Studies of whisking behavior in freely behaving animals reveal two main whisking modalities. Away from an object, rodents perform “exploratory” behavior characterized by extended whisking protractions, allowing the animal to scout larger areas. On the other hand, when they are in contact with an object, they adopt a “scanning” strategy, reducing the amplitude of whisking protractions to match their distance to objects.

Matching whisking protraction to object distance is an effective strategy to ensure that contacts happen at the end of the protraction, with a light touch and minimal bending - a strategy called “minimal impingement”. After a contact with an unexpected object, whiskers cease to protract very rapidly (e.g., within 15ms). This rapid cessation of protraction (RCP) suggests a rapid feedback-control mechanism [9, 10, 11].

However, feedback-control alone cannot account for the fact that when objects are removed, the target protraction remains stable for at least one whisking cycle, increasing again only afterwards [7]. This finding suggests that rodents target their whisker protractions to where they *expect* objects to be, rather than just react to unexpected contacts. In other words, the modulation of whiskers’ amplitude appears to be an anticipatory strategy that depends on sensory prediction errors - or the mismatch between expected and sensed inputs - rather than just on current sensory inputs.

Importantly, the anticipatory modulation of whiskers allows the animal to actively perceive - and maintain an expectation about - its distance from an object. Here, perception is “active” in the sense that the alignment between the animal’s expectations about its distance from an object and the effective distance is achieved by acting (i.e., by changing the amplitude of whiskers’ oscillations), not by merely updating internal beliefs, as normally assumed in inferential theories of perception [12, 13, 14, 15]. Unlike classical theories of perception, which assume that the brain recognizes objects via representations of (hiearchies of) action-independent object features [16, 17], this form of active perception relies on a *generative* model that describes how (touch) sensations change when an object is in contact, i.e., models of sensorimotor contingencies [18] (see [19, 20] for examples of such generative models).

In this paper, we formally characterize rodents’ anticipatory and error-correction whisking strategies in terms of active inference: a framework developed in computational neuroscience to explain animal behavior and neural activity as resulting from the minimization of variational free energy, or under simplifying assumptions, prediction errors [21]. We develop an active inference model of whisking dynamics and use it to simulate an object localization task that exposes anticipatory aspects of whisking dynamics [7]. Furthermore, we discuss how our simulated agent performs active perception, displaying a dynamical alignment of internal expectations and animal-object distance without explicitly encoding such distance. Finally, we illustrate the putative neurobiological substrate of the active inference model of rodent whisking.

### 1.1. Summary of the active inference perspective on active whisking

The core idea of the active inference approach proposed here is to specify a generative model of sensorimotor contingencies: contingencies between whiskers’ protractions and the resulting proprioceptive and somatosensory (touch) sensations.

The agent’s model continuously generates proprioceptive and touch-related predictions during the action-perception loop. Crucially, the model includes a prior belief that objects will only be touched at the end of a whisker protraction. Hence, the agent will continuously adjust (increase or decrease) whisker protractions to match the prior prediction - or in other words, to minimize prediction errors resulting from either touching unexpected objects, or failing to touch expected objects that are instead missing.

The dynamical adjustment of whiskers’ protractions automatically produces a transition from exploration to scanning with a decrease of whisking amplitude, and from scanning to exploration with an increase of whisking amplitude. Importantly, both the decrease and the increase of whiskers’ amplitude result from the same (error minimization) mechanism, in two different conditions.

During the exploratory phase (characterized by larger oscillations of the whiskers), touching an object with the whiskers before the end of the protraction generates a *somatosensory (touch) prediction error*. This is because such input is unexpected given the prior (that an object will be touched only when the whiskers are fully extended). To minimize this prediction error, the animal can *reduce* the amplitude of the whisking protractions, so that it matches more closely the distance from the object. At the same time, the animal can also update its prior belief about the whisking amplitude, so to select (or predict) a smaller amplitude for the next step. This dynamical adjustment of whisking amplitude entails a shift from an “exploratory” to a “scanning” phase.

During the object scanning phase, when somatosensory or “exploratory” prediction errors have been minimized and hence the amplitude of whisker protractions closely matches animal-object distance, the sudden removal of the object causes a new sensory prediction error to arise - because the animal expected a touch sensation in correspondence of the missing object. To minimize this new prediction error, the animal will thus *increase* the amplitude of the whisking protraction, until another object is located, or the maximum span of the protractions is reached. In parallel, it will also update its prior beliefs about these reaching movements, leading to a larger protraction amplitude for the ensuing cycles. This dynamical adjustment of whisking amplitude characterizes a transition from a “scanning” to an “exploratory” phase.

As we will see in our results, in addition to producing switches from exploration to scanning and vice versa, the dynamic adjustment of whisker movements allows an animal to perceive its distance from an object. Indeed, in a regime of dynamical convergence (i.e., after some cycles of adjustments) the whiskers’ amplitude can be used as a proxy for animal-object distance – even if the generative model has no explicit notion of either objects or distance, but simply embodies sensorimotor contingencies, or relations, about how whisker protractions generate touch sensations. This idea is in accordance with theoretical views of perception as a closed-loop convergence process, during which motor variables (e.g., whisking velocity or amplitude) are dynamically controlled over time, until they converge to steady state [22, 6, 23].

In the next section, we formally introduce the active inference model and illustrate simulations of whisking behavior during animal-object interactions [7]. Our simulations will show that minimizing somatosensory prediction errors guides both active exploration and the scanning of objects by whiskers.

## 2. Methods

### 2.1. Active inference

In theories of agency and perception inspired by Bayesian principles, the goal of an inferential agent is to generate a purposeful engagement with the world via the estimation of the probability of hidden variables **u** = {**x, v**}^1^ given by hidden states 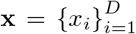 and hidden causes 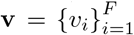 for some sensory inputs 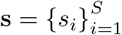

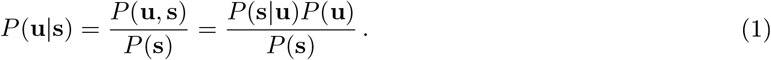

A common issue with exact Bayesian schemes is that the marginal likelihood or model evidence *P*(**s**) is often analytically intractable or computationally difficult to calculate. Moreover, the posterior *P*(**u**|**s**) may not follow a standard distribution and thus have no tractable summary statistics.

Active inference [25, 21, 26, 27, 28] brings forward a biologically plausible (variational) approximation to this problem, where an auxiliary distribution *Q*(**u**) called *recognition density* has to be optimized to become a good approximation of the posterior. To implement this, the Kullback-Leibler divergence (a measure of the dissimilarity between probability distributions with roots in information theory) is minimized:

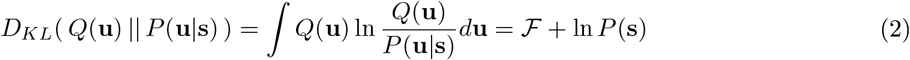

where ℱ ≡ − ⟨ln*P*(**s, u**) ⟩_*Q*_ + ⟨ln*Q*(u) ⟩_*Q*_ is the *variational free energy* (VFE). This quantity depends on the recognition density and the agent’s knowledge about the environment recapitulated by the joint density *P*(**s**, **u**) = *P*(**s**|**u**)*P*(**u**), while ln*P*(**s**) does not depend on the recognition density *Q*(**u**). Hence, minimizing ℱ with respect to *Q*(**u**) will minimize the whole *D*_*KL*_.

Optimizing ℱ for arbitrary *Q*(**u**) can however be a complex task. A common choice is thus to introduce a Laplace approximation [29] assuming that the joint density is a smooth function of **u**, and that its logarithm can be approximated with a quadratic function near the mode (i.e. the logarithm of a Gaussian distribution). This is equivalent to a variational Gaussian approximation of the recognition density as a multivariate Gaussian form over the D-dimensional space **u** centered around the estimated mode ***μ*** [30]. The Laplace approximation is therefore here used to evaluate, up to quantities that act as constants during the minimization process, ℱ as

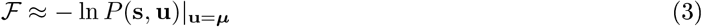

where ***μ*** has the role of the agent’s expectations about the hidden states **x** and hidden causes **v** (see [31] for a pedagogical treatment).

To evaluate the VFE, we then specify the joint density *P*(**s, u**) expressing hidden states and sensory inputs. In particular, we will assume that

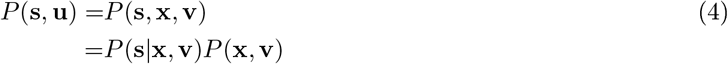

thus specifying dependencies typical of state space model in continuous time, with the likelihood *P*(**s**|**x, v**) corresponding to observation law and the prior *P*(**x, v**) encoding dynamics following the treatment found, for instance, in [24]. At the same time, we also assume that the probability of hidden causes **v, *P***(**v**), is Gaussian with precision approaching zero (i.e., whose covariance tends to infinity), meaning that it will not affect the remainder of the prior *P* (**x**|**v**). In the case of a system with one hidden state, one hidden cause and one sensory input this is equivalent to

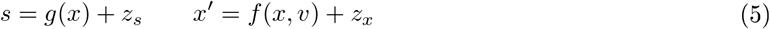

where *g*(·) maps hidden states to observations, *f* (·) encodes the dynamics of the hidden state and *z*_*s*_,*z*_*x*_ represent (white) noise terms. Under zero-mean Gaussian assumptions about random variables *z*_*s*_ and *z*_*x*_, the VFE is then

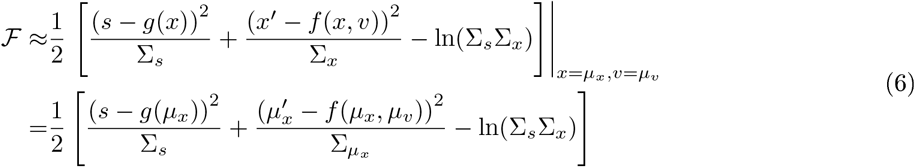

where ∑_*s*_ and ∑_*x*_ are the variances of the random variables *z*_*s*_ and *z*_*x*_, respectively.

At this point, the optimization of the VFE can be achieved either by changing the expected states *μ, μ ′* through a (modified) gradient descent (see [24] for details)

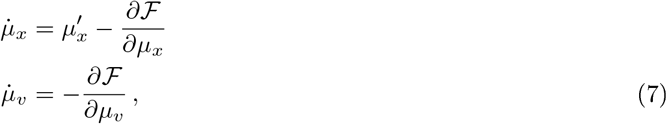

or by selecting an action through a free energy gradient descent where

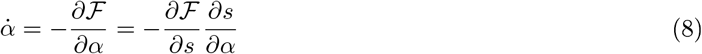

The gradient with respect to actions highlights a key assumption of active inference treatments, whereby the action variable *a* is not itself part of the generative model (to be specified in the next section) but instead appears only in the recognition dynamics (i.e., the minimisation of variational free energy for action and perception embodied by an agent) via the relation 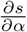, capturing how actions *a* affect sensory input *s* given an agent’s specific “implementation”, e.g., bodily constraints. In the active inference literature this is usually associated with the idea of reflex arcs, proposed to be the central construct for motor execution (here thought of essentially as a chain of reflexes) via the minimization of proprioceptive prediction errors induced by top-down modulatory signals from motor areas [32]. Such signals (i.e., predictions about proprioceptive sensory input) are thought to generate mismatches between *expected* proprioceptive signals and effective proprioceptive sensations, based on an equilibrium-trajectory (referent) control [33, 34] framework encoding desired movements in the (low-dimensional) space of observables, as opposed to classical treatments of motor control relying on inverse models that specify motor commands as (high-dimensional) signals in an intrinsic (bodily) frame of reference describing movements explicitly (e.g., motor signals describing stretching and compressing of muscle fibers).

### 2.2. Active whisking model

Whisker control in rodents is simulated in a rat-like agent that controls a single whisker, attached near its nose, in a 2D kinematic environment (see Figure 1A). The agent moves forward with a constant speed until it reaches a specified final position (e.g., the end of a platform). At that point, an object can appear within reach of the whisker, and later disappear. Our simulations (discussed in Section 3) will show how the agent controls its whiskers’ protraction when objects appear or disappear.

**Figure 1:**
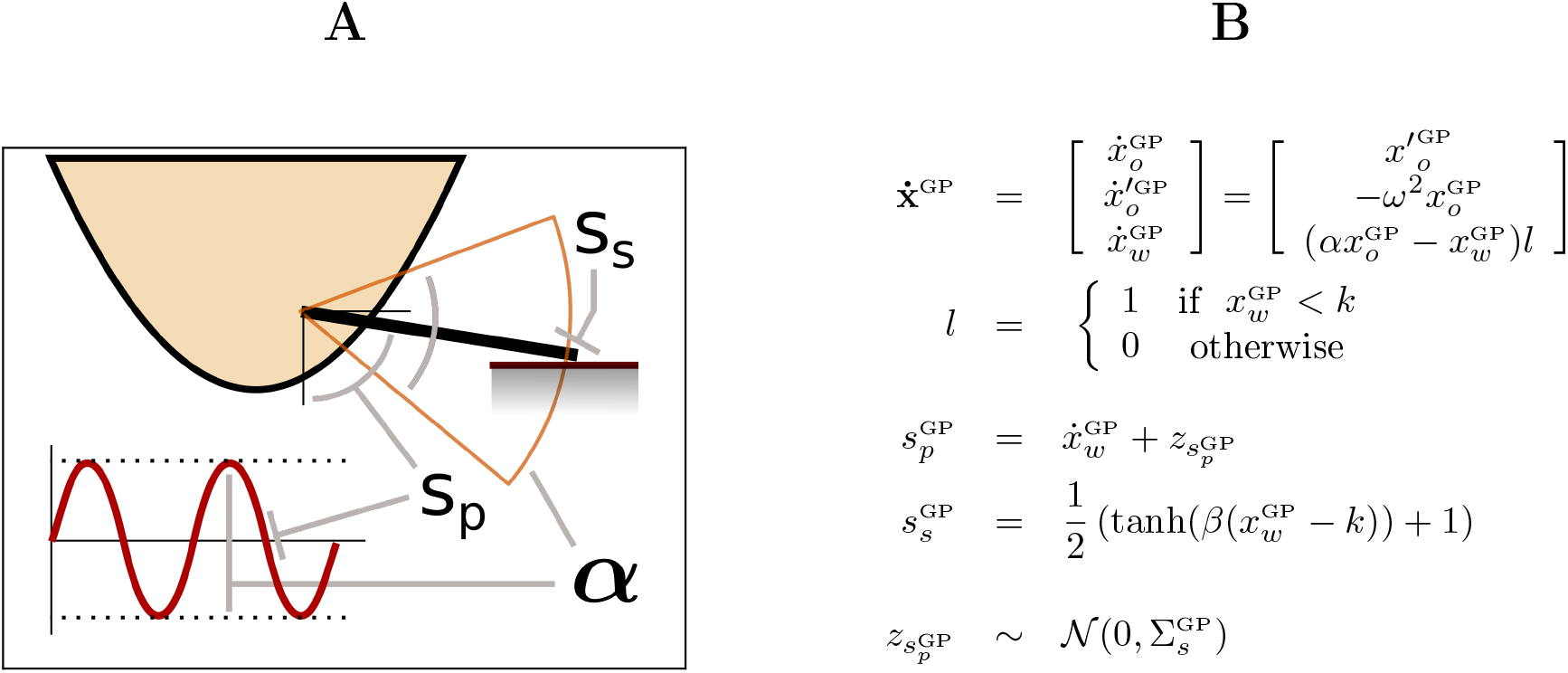
The simulated environment, or generative process. (A) The 2D environment: the simulation includes a rat-like agent with a single whisker. (B) Generative process and whisker dynamics. The generative process is specified by the dynamics of the state variable **x**^GP^. The first two components of **x**^GP^(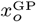 and 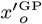) represent a central pattern generator described by an harmonic oscillator, while the last component 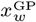 describes the motor command for the angular position of the whisker’s joint. The parameter *ω* determines the frequency of the central pattern generator oscillation and is kept constant during all simulations. The value of *α* specifies a scaling factor for the oscillation in the 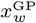 component. The generated sensory states are defined by 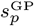 and 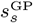, with *k* representing the current upper limit of 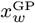 (the position of the external object in terms of whisker base angle), and *β =* 500 being the smoothness of the pressure response of the whisker.

Importantly, in active inference, there is a fundamental distinction between the *generative process*, corresponding to the actual dynamics that govern the whisker’s movements in the environment (Figure 1) and the agent’s (whisker) controller, implicitly specified as a *generative model* of the whisker’s dynamics (Figure 2) giving rise to the agent’s recognition dynamics (see previous section). The recognition dynamics and generative process are thus engaged in a closed action-perception loop: the recognition dynamics instantiate actions for the generative process and the generative process provides sensory evidence to the recognition dynamics. To set up such action-perception loop, we will describe next the generative process and the generative model used to derive the recognition dynamics of the agent, respectively.

**Figure 2:**
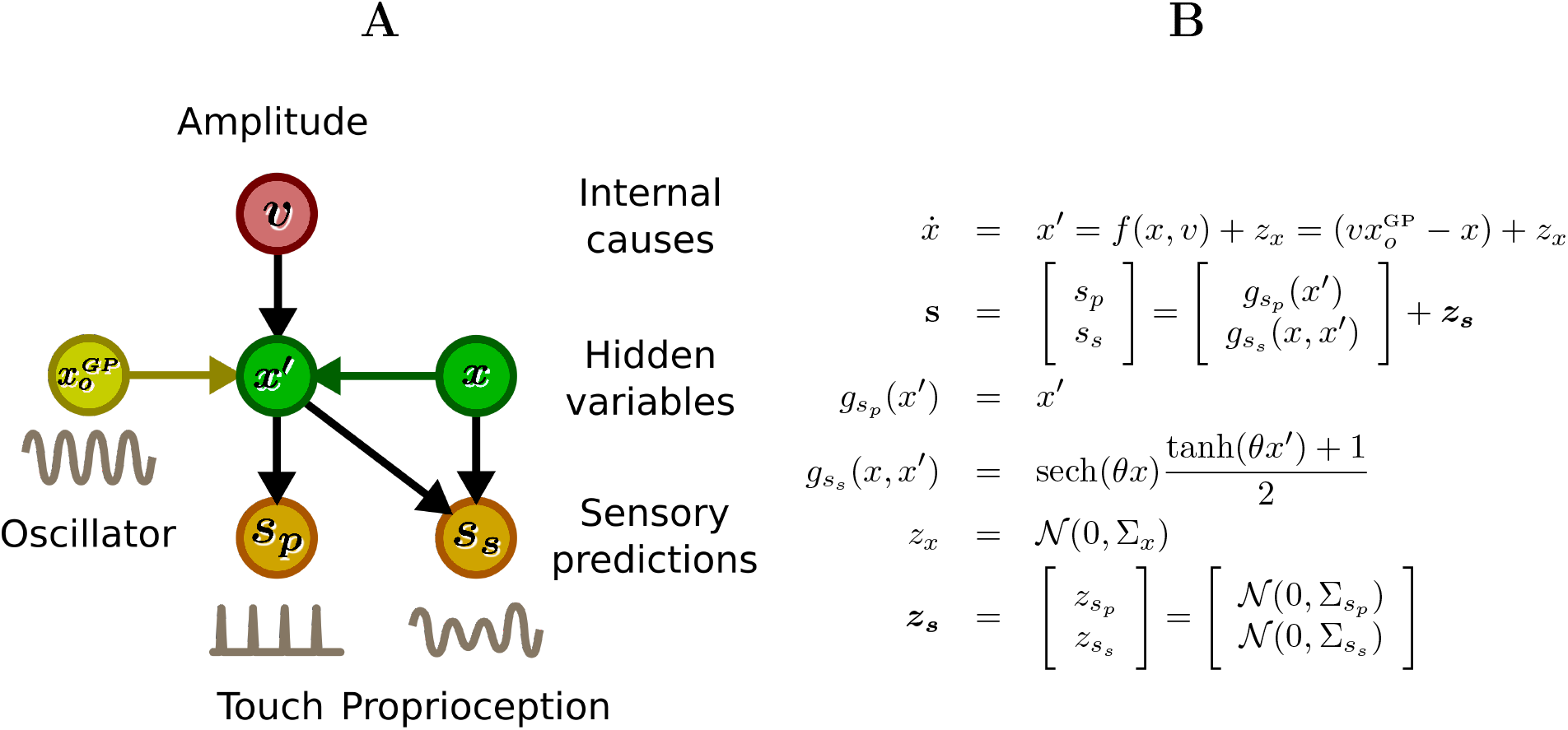
The generative model. (A) Schematic illustration of the model as a Bayesian network. The model includes *x*, a variable that filters the amplitude of the whisker oscillation given by the central pattern generator 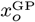. The variable *x* depends also on an input (*υ*), encoding a prior over the desired amplitude. *x* and *x′* are then mapped to predictions about somatosensory (*s*_*s*_) and proprioceptive (*s*_*p*_) input. (B) Generative model in analytical form. The somatosensory mapping function 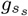 between *μ*_*x*_ and *s*_*s*_ has *θ* = 50 that sets the smoothness of the pressure sensory prediction. The yellow arrow from 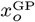 describes a known input to the internal variable *x′*, under the assumption of an additional observation with an identity mapping to the pattern generator 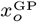, omitted from the figure for simplicity (i.e., identity mappings imply prediction errors with infinite precision, or equivalently, zero variance, and thus variational updates instantaneously reaching steady-state).

#### Generative process

The whisker’s dynamics are simulated within the generative process (Figure 1B). The whiskers’ oscillatory movements are given by a dynamical system which comprises of three variables. The first two (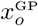 and 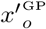) define a harmonic oscillator of unitary amplitude as a first order system, which (for simplicity) will remain fixed in our simulations. The third one 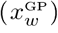 defines the actual movement of the joint angle of the whisker. This auxiliary variable rescales (with a leak term) the amplitude of the oscillator response in 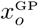 through a parameter *α*. Importantly, in this model, the agent’s actions correspond to updates of *α*, the parameter that regulates the amplitude of the oscillation of 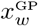. In other words, in this model, the agent can only change the way it samples its environment, by modulating the amplitude of the protraction of its whiskers.

The term 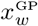 is not only controlled by the agent, but also constrained by an external factor (*k*) that corresponds to the current limit of the whisker angle, due for example to the presence of an object that limits its oscillation. When the whisker angle (provided by 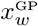) approaches the limit *k*, a somatosensory touch event is produced, which corresponds to a peak in the sensory state 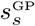 of the agent’s generative model. Proprioceptive sensory states are simulated as linear and noisy mappings of 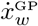.

#### Generative model

The generative model is schematically illustrated in Figure 2. For simplicity, this model only describes only the dynamics of a single whisker; the forward movement of the simulated agent is hardwired in our simulations and hence not included in the generative model. Here, the variable 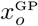 encodes the central pattern generator of the generative process. It is assumed to be known (analogously to standard problems of filtering with known inputs [35], and in line with dynamical accounts of sensorimotor interactions with no fixed set-point control, cf. the role of inputs *φ*_*0*_ in the centrifugal governor found in [36]) and thus equivalent to a sharply peaked Gaussian distribution with (in the limit) zero variance.

The variable *x* represents the (stochastic) dynamics of the whisker and is described in Langevin form, with a deterministic term rescaling (with a leak) the amplitude of the oscillations of 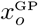 by a factor *ν*. Here we also implicitly assumed the presence of an additional observation variable with an identity mapping (i.e., infinite precision) to the pattern generator 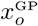. Due to the mapping appearing as a simple identity, we however omitted this from the generative model and thus from the associated free energy functional.

On the other hand, the hidden cause *υ* models an exogenous input (from a dynamical systems perspective) in the dynamics of *x*, scaling the amplitude of whisker oscillations while influenced by bottom-up prediction errors, see below. Crucially, *P*(*υ*) is modelled as a Gaussian distribution with arbitrarily high variance (zero precision), meaning that its role as prior on the desired amplitude is entirely driven by external stimuli, i.e., *υ*, or rather its mode *μ*_*υ*_, will act a proxy that registers the presence of unknown objects in the environment.

The variable **s** includes two sensory predictions: a proprioceptive prediction (*s*_*p*_), proportional to the current angle velocity of the whisker, and a somatosensory (touch) prediction (*s*_*s*_), corresponding to the actual pressure over the whisker during the contact with an external object. Both predictions are assumed to be generated from Gaussian distributions, with means given by the two mapping functions 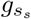 and 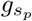 that link the dynamics of the variable *x*. The prediction *s*_*s*_ is modelled as a radial function centered on the local maximum of the hidden states encoding the whisker’s angle (i.e. the component *x* of the internal state variable and its velocity *x*′). It is then interesting to note that while *s*_*s*_ and 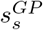 refer to the same physical quantity, there is a fundamental difference between the radial function used to model *s*_*s*_ and the equation used in the generative process for 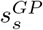 (see Figure 1), see also [19, 20].

#### Variational updates, or recognition dynamics

As explained in Section 2.1, once the joint density of the generative model (Equation 4) is specified, it is possible to compute the variational free energy using the current means (modes) of the model’s hidden variables 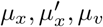 and the current sensory states (see equation 6). With the VFE specific to the generative model, we can then find the gradients of VFE, generating updates for perception and action of our simulated agent (see equations 7 and 8). For perception, the model is updated by changing the posterior beliefs about both the expected hidden states *μ*_*x*_ and 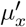 and the expected causal variable *ν*. For action, the model is updated by descending the free energy gradient 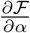 with respect to the parameter *α*. This gradient can be factored into two components 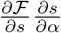. The former 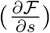 is the gradient of the free energy with respect to the sensory states. The latter 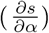 is the gradient of sensory states with respect to the action. It corresponds to a learned (or hardwired) mapping between action and sensory states at the level of the generative process and it is usually identified with the spinal neural circuits which manage reflex arcs [37]. This gradient 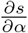 is implemented here as a composition of 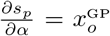 and 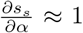, where 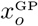 is the exact solution of the proprioceptive part of the gradient 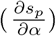, while the somatosensory gradient 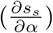 is approximated to 1, which is sufficient to inform the process about the direction of the update of *α* based on somatosensory states.

The update of *ν* showcases how the agent adapts to external constraints, such as the presence of an object that blocks whisker movements, updating its beliefs about the causes of its sensory input and preventing whisker oscillations from reaching the desired, but unattainable, amplitude specified in the current prior expectation *μ*_*ν*_. In other words, when the agent cannot change the world (e.g., cannot use the whisker to push the object) via its own actions, it can nonetheless adapt its internal prior beliefs *μ*_*ν*_ to account for the actual constraints of the generative process.

## 3. Results

We used the active inference model of whisking to simulate the experimental setup of Figure 1a, and the control of a single whisker in the presence or the absence of an object. The simulation can be conceptually divided into three phases. (1) Initially there is no object and the animal’s whiskers operate in “exploration mode”, a phase characterized by large whisker oscillations. (2) An object then appears and the animal progressively reduces whisking amplitude, shifting from “exploration” to the “scanning” of the object. (3) Finally, the object is removed and the amplitude progressively increases once again, marking a transition back from “scanning” to “exploration”.

Figure 3 shows the results of the simulations. At first, the recognition dynamics and the generative process are in the same initial conditions. Figure 3A shows the dynamics of the variable 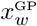 that controls the amplitude of the whisker oscillation in the generative process, as a function of time. Without objects, the animal is in “exploration” mode, characterized by large whisker oscillations and extended protractions, which slightly increase over time. In the second phase, an object (black line) appears, limiting whiskers’ protractions. Finally, in the third phase, the object disappears and whisker protractions increase once again. Crucially however, they initially remain significantly smaller than in the first phase. This is especially evident in the first cycle after the disappearance of the object, when they almost match (expected) animal-object distance, even if the object is not there to stop whiskers’ protractions. This is the key empirical observation that has lead to the proposal that whisker movements are guided by anticipatory mechanisms that consider expected animal-object distance, rather than feedback-control mechanisms [7].

**Figure 3:**
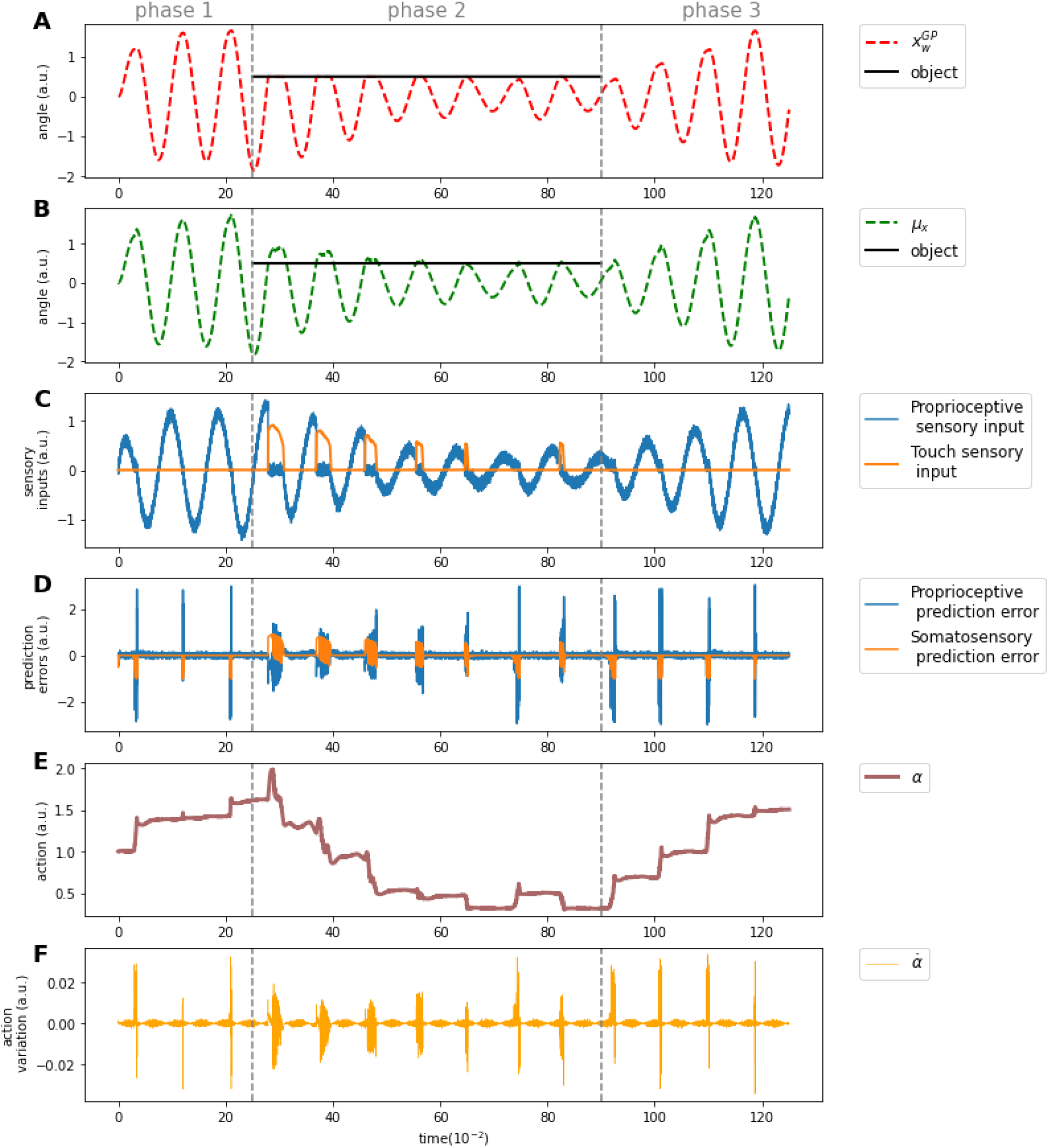
Simulation results. (A) Dynamics of the variable 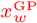, which corresponds to the whisker’s base angle in the generative process. (B) Dynamics of the variable *μ*_*x*_, which corresponds to the predicted whisker’s base angle in the generative model. By comparing A and B, it is possible to notice that the appearance of the object in phase 2 affects immediately the dynamics of the whisker’s base angle in the generative process 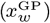; however, the dynamics of the predicted whisker’s base angle *μ*_*x*_ take slightly longer to adapt. (C) Agent’s sensory inputs (somatosensory (touch) in orange and proprioceptive in blue). (D) Agent’s prediction errors (somatosensory (touch) in orange and proprioceptive in blue). (E-F) Dynamics of *α* (action) and its derivative 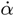. Negative peaks of touch prediction error correspond to situations in which the agent would have expected to touch the object but did not - and hence increases the amplitude of the whisker protraction (by increasing *α*). Rather, negative peaks of touch prediction error correspond to situations in which the agent feels unexpected touch sensations - and hence decreases the amplitude of the whisker protraction (by decreasing *α*).

In the first phase, when there is no object, the variables 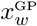 and *x* that control the amplitude of whisker oscillation in the generative process (Figure 3A) and model (Figure 3B) slightly increase over time. This is because the agent expects to make contact with something at the end of the whisker protraction, but does not receive any touch input (Figure 3C) – and hence generates a negative somatosensory prediction error (Figure 3D). The action increment that cancels out this prediction error (Figure 3F) increases the value of *α* (Figure 3E), which in turn amplifies the whisker oscillation / protraction (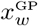in Figure 3A).

In the second phase, when the object (black line) appears, the agent’s expected amplitude is significantly greater than the actual amplitude of the generative process; this is because 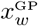 is physically blocked by the presence of an unexpected object, but *μ*_*x*_ is not (note also that there is no explicit representation of the object in the generative model). However, the variable *μ*_*x*_ rapidly re-aligns to the true dynamics of the generative process. This re-alignment process is guided by an error-correction mechanism, as we can see with sensory signals and prediction errors shown in Figure 3C and Figure 3D, respectively. As shown in the third panel, the agent receives somatosensory (touch) and proprioceptive sensations from the generative process, especially evident with the first prominent somatosensory input (first orange peak). This input is completely unexpected and hence generates a large positive somatosensory prediction error, which is shown in Figure 3D. The (negative) action increment that cancels out this prediction error (Figure 3F) suddenly decreases the value of the *α* parameter (Figure 3E), which in turn decreases the amplitude of the whisker oscillation / protraction (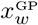 in Figure 3A). This decrease stops when the whisker touches the object gently and the *α* parameter finds a dynamic equilibrium around the correct value.

Finally, in the third phase, when the object (black line) disappears, the *μ*_*x*_ variable (Figure 3B) continues to guide whisker protraction at the expected distance from the object, at least in the first cycle. The sudden disappearance of the object implies that the agent does not receive its expected touch sensation (i.e., there is no orange peak in Figure 3C) and hence generates a negative somatosensory prediction error (Figure 3D), which is compensated by progressively increasing the amplitude of whisker oscillations (Figure 3E and Figure 3F). This marks thus a transition back to the exploratory mode characterising phase one, bringing the whisker oscillations back to baseline, i.e., extended protactions to explore the environment.

The simulation of the three phases closely resembles the empirical results reported in [7], where upon appearance of an object (in [7], a platform), whiskers’ protractions match precisely the distance between the animal and the object – as seen in the second phase of our simulation. Crucially, when the object is suddenly removed, whiskers’ protractions do not adapt immediately, but remain proportional to the last (expected) animal-object distance – which is exactly what happens in the third phase of our simulation. This suggests that the control of whiskers may be anticipatory at its core (based on expectations about animal-object distance) [7]. Furthermore, our simulations show that error-correction (via a free energy minimization) mechanism can explain both the shift from “exploration” to “scanning” when the object is introduced (second phase) and the reverse one from “scanning” to “exploration” when the object is removed (third phase). In our model, what guides the changes in whisker protraction is the somatosensory (touch) prediction error. The proprioceptive prediction error instead plays a stabilizing role on whisking oscillation, allowing our agent to compensate for small and unexpected changes caused by external disturbances of noise.

### 3.1. Control of multiple whiskers

To showcase further applications of our model, we finally consider an agent with multiple (here, two) whiskers. Given the structure of our framework, the formulation is rather straightforward, showing thus its potential for the development of more complex models of active perception. In this extended generative model (Figure 4A), we simply add a copy of the variables specified in the previous generative model, except for the oscillator variable 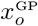, which is shared across all the whiskers. Given this duplication, the parameters of different whiskers can be tuned independently, hence we here simulate two whiskers with slightly different lengths and centers of oscillations (Figure 4). However, since the model relies on a shared oscillator variable 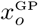, the resulting whisker dynamics remain in phase during scanning behaviour (Figure 4B) – as observed empirically [6].

**Figure 4:**
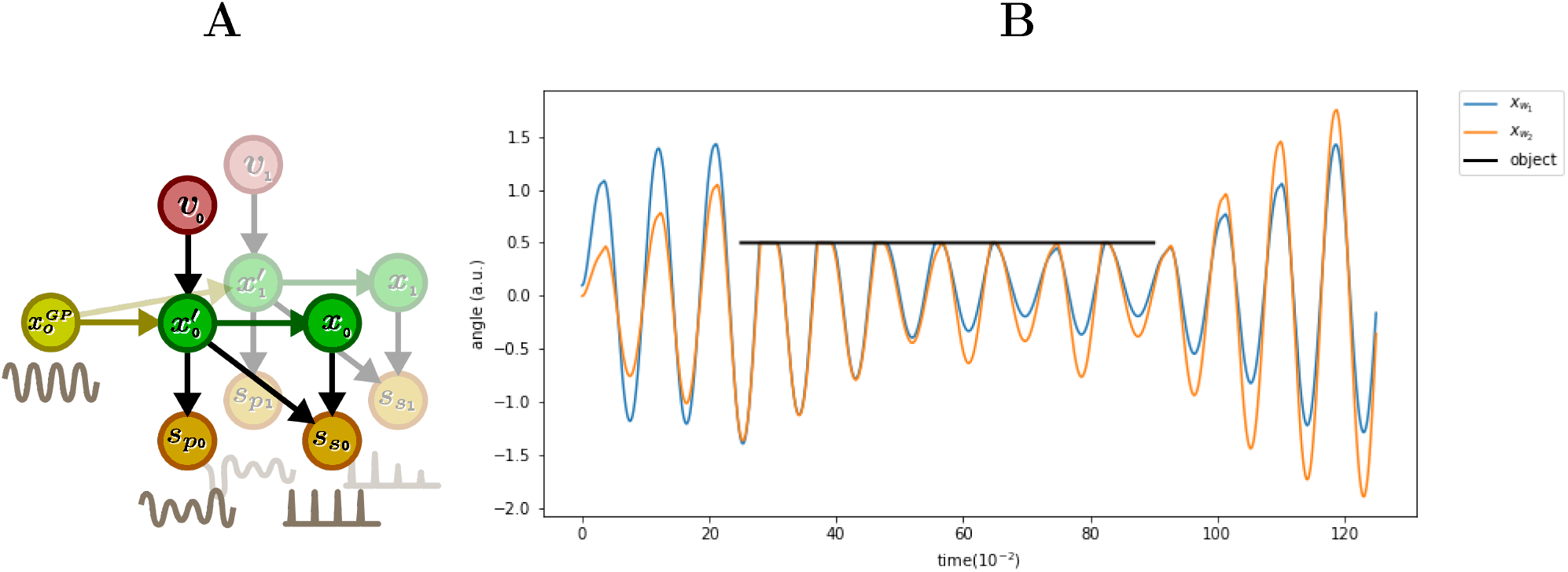
Simulation of the control of two whiskers having slightly different dimensions and central points of oscillations. (A) Extended generative model for the control of two whiskers. The model’s variables are the same as Figure 2, but there is a separate set of variables for each whisker, except for the oscillator variable 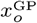, which is shared across both whiskers. (B) Simulation results. The notation is the same as in Figure 3. Here, they key thing to notice is that the two whiskers (blue and orange) have oscillate in phase. See the main text for an explanation.

## 4. Discussion

We proposed an active inference model of active sensing through whiskers. We started from the premise that rodents use their whiskers’ dynamics to make (perceptual) inferences about the world; and described the active control of whiskers as an error-correction mechanism ensuring that a prior prediction (that something will be touched at the end of the protraction) can be realized in the environment.

Our simulations illustrate that the active inference model reproduces key anticipatory aspects of whisker movement control, such as the fact that mice target the amplitude of their whisking movements to the expected distance from an object; and (contrary to purely reactive models) that such expectations are robust and stable for some time even when objects disappear [7]. Furthermore, the model characterizes the the alternation of exploration and scanning phases, exploration to scanning when an object is sensed and scanning to exploration when such object disappears, in terms of a unique prediction error minimization mechanism. Finally, this model explains how the control of whisker movements supports active perception by dynamically aligning expectations (about whisker protraction) in the agent’s generative model to effective object distance. This is because the same error (or more precisely, free energy) minimization process has the dual role of selecting the motor actions to execute and updating the internal variables of the generative model.

The proposed model builds on a tradition of early cybernetic models and in particular perceptual control theory [38], which stresses the importance of continuously controlling motor variables (e.g. modulating the force impressed on the acceleration pedal while driving) to keep a preferred (prior) perceptual variable constant (e.g., to ensure that I always see 120Km/h on my speedometer). Furthermore, our model is also in line with the (more recent) theoretical view of active perception as a closed-loop convergence process, where motor variables (here, whisker amplitude) are continuously modulated for perceptual purposes [22, 23].

The active view of perception that emerges from the model differs in at least two ways from views of perception as the recognition of objective features of the environment [16] or the inference of external variables [12, 13, 14, 15]. First, traditional theories of perception assume that in order to recognize external variables (e.g., objects, faces or places), an agent has to be endowed with a model of these variables and their features. Rather, the proposed model stresses the perception of external variables (here, animal-object distance) without an explicit, action-independent model of objects and their features. The generative model only includes knowledge about touch sensations changing as an effect of whisking actions (i.e., sensorimotor contingencies [18, 39, 19, 40]), without explicit action-independent perceptual features or external variables like objects or distances, as commonly assumed in other theories of perception.

Second, traditional theories assume that perception is only achieved by changing beliefs internally. Here, instead, the perception of animal-object distance is achieved in an active manner: by dynamically adjusting a motor variable (whisker amplitude) based on a prior encoding the sensorimotor coupling between agent and environment, rather then a fixed set-point [36]. This ensures that touch sensations only happen at the end of the protraction - hence minimizing any discrepancy between prior predictions about touch and current sensations. An interesting “side-effect” of this adjustment is that it provides an implicit estimate of animal-object distance. Given this lawful correlation between whisker protraction (in the stationary regime) and animal-object distance, the animal can use the former as a proxy for the latter, so to, for example, approach or avoid the object. Our model therefore shows how to endow active inference models with minimalistic (sensorimotor or action-oriented) models or action-world interactions, in place of more complete models that describe and predict external dynamics independent of an agent’s actions [41, 42, 43, 44, 45, 46, 47] (see also e.g., [19, 48, 36, 20] for example implementations). An objective for future research is testing to what extent this approach scales up to robotic settings [49, 50].

### 4.1. The neurobiology of whisking from an active inference perspective

The neurobiology of active whisking involves a widely distributed brain network, which spans sensory, motor, premotor/prefrontal areas and the brain stem [1]. Figure 5 shows part of this network and highlights the putative neuronal underpinnings of the main variables of the generative model used in this article. We map sensory inputs and prediction errors in the two sensory modalities (somatosensory information about touch, such as the mechanical pressure over whiskers *s*_*s*_ and proprioceptive information *s*_*p*_) to the barrel cortex (vibrissal somatosensory cortex) [51, 52, 53], the two variables of the central pattern generator (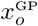 and 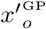) to the brain stem, the hidden variable that modulates the central pattern generator *μ*_*x*_ to vibrissal motor cortex and the hidden cause *μ*_*ν*_to the premotor / prefrontal cortex.

**Figure 5:**
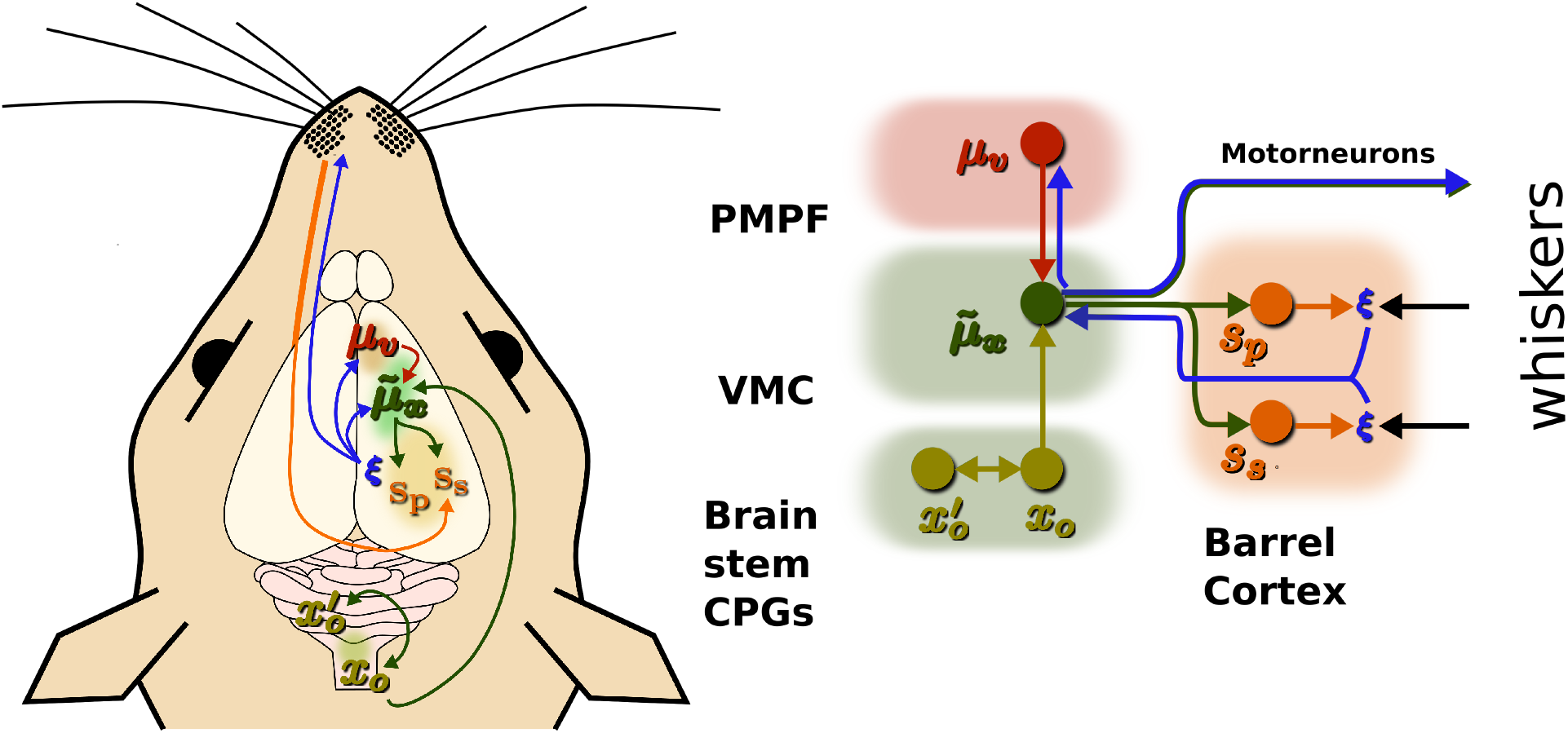
Putative neuronal underpinnings of the generative model for whisker control discussed in this article. This simplified neuronal scheme comprises premotor / prefrontal areas (PMPF), vibrissal motor cortex (VMC), central pattern generators (CPGs) in the brain stem, and Barrel Cortex / vibrissal somatosensory cortex. Colored circles represent variables of the generative model, whereas the *ξ* symbols represent prediction errors. Colored edges show the (neuronal) message passing between variables of the model. The red edge originating from the *ν* (causal) variable represents top-down signals. Blue edges originating from the *ξ* symbols represent prediction errors that are propagated bottom-up. See the main text for explanation.

The proposed neurobiological scheme follows a dynamical systems perspective on cortical computations and movement control [54, 55]. In this scheme, premotor/prefrontal areas define a prior over the amplitude of the oscillation of whiskers. Vibrissal motor cortex controls whisker movements (until they reach the desired set point) by modulating the amplitude and midpoint of whiskers protractions [56, 57]. The actual control of movement engages synergistically the vibrissal motor cortex and central pattern generators in the brain stem. Evidence indicates that vibrissal motor units indirectly control the activity of vibrissal motoneurons, through the modulation of sub-cortical central pattern generators; and in turn the motorneurons control the facial muscles responsible for whiskers’ movements [58].

Somatosensory prediction errors (discrepancies between somatosensory predictions and sensations) are computed by the barrel cortex and are related to whisking amplitude change rather than whisking frequency or velocity [59, 7]. These prediction errors are sent to vibrissal motor cortex and premotor/prefrontal cortical areas, to keep the agent’s generative model in register with external constraints, such as the (unpredicted) presence of an object, which may prevent the whisker from reaching the “desired” amplitude specified in the current prior expectation (*μ*_*ν*_). Specifically, prediction errors modify the oscillation amplitude of whiskers at the level of motor cortex; and the prior about oscillation amplitude at the level of premotor/prefrontal cortical areas. Finally, the model includes a second type of (proprioceptive) prediction error, which measures the difference between the expected and currently sensed whisker protraction, and ultimately helps stabilizing whisker movements in face of small external disturbances, such as wind.

## 5. Conclusions

The model introduced in this article explains the active control of whisker movements as an active inference process which continuously minimizes prediction errors and the discrepancy between expected somatosensory sensations and current observations. This anticipatory, error correction mechanisms allows smooth transitions between phases of exploration and scanning of objects, and vice versa, empowering the animal with the ability to (actively) perceive their distance from objects.

Our model is in agreement with key empirical evidence supporting the anticipatory, expectation-based nature of whisking control [7]. Furthermore, it provides preliminary evidence for a new interpretation of neural signals in the brain circuits for whisking in terms of Bayesian (active) inference constructs, such as priors about amplitude of oscillation in premotor / prefrontal cortices and sensory prediction errors in barrel cortex. While some aspects of the model qualitatively align with the known neurophysiology of whisking (see Section 4.1) the specific mappings between neuronal circuits and active inference signals will be an object of future studies.

## Acknowledgments

This research has received funding from the European Union’s Horizon 2020 Framework Programme for Research and Innovation under the Specific Grant Agreement No. 945539 (Human Brain Project SGA3) and the European Research Council under the Grant Agreement No. 820213 (ThinkAhead). MB is a JSPS Postdoctoral Research Fellow supported by a KAKENHI Grant-in-Aid for Scientific Research (No. 19F19809).

## Data Availability

The python code to reproduce the results of this paper can be found at: https://github.com/francesco-mannella/Active-Inference-In-Continuous-Time-With-Whiskers

## Footnotes

1 For a treatment including other variables, e.g., parameters evolving on different time scales, see for instance [24].

